# A modified sequence capture approach allowing standard and methylation analyses of the same enriched genomic DNA sample

**DOI:** 10.1101/209585

**Authors:** Lisa Olohan, Laura-Jayne Gardiner, Anita Lucaci, Burkhard Steuernagel, Brande Wulff, John Kenny, Neil Hall, Anthony Hall

## Abstract

**Background:** Bread wheat has a large complex genome that makes whole genome resequencing costly. Therefore, genome complexity reduction techniques such as sequence capture make re-sequencing cost effective. With a high-quality draft wheat genome now available it is possible to design capture probe sets and to use them to accurately genotype and anchor SNPs to the genome. Furthermore, in addition to genetic variation, epigenetic variation provides a source of natural variation contributing to changes in gene expression and phenotype that can be profiled at the base pair level using sequence capture coupled with bisulphite treatment. Here, we present a new 12 Mbp wheat capture probe set, that allows both the profiling of genotype and methylation from the same DNA sample. Furthermore, we present a method, based on Agilent SureSelect Methyl-Seq, that will use a single capture assay as a starting point to allow both DNA sequencing and methyl-seq.

**Results:** Our method uses a single capture assay that is sequentially split and used for both DNA sequencing and methyl-seq. The resultant genotype and epi-type data is highly comparable in terms of coverage and SNP/methylation site identification to that generated from separate captures for DNA sequencing and methyl-seq. Furthermore, by defining SNP frequencies in a diverse landrace from the Watkins collection we highlight the importance of having genotype data to prevent false positive methylation calls. Finally, we present the design of a new 12 Mbp wheat capture and demonstrate its successful application to re-sequence wheat.

**Conclusion:** We present a cost-effective method for performing both DNA sequencing and methyl-seq from a single capture reaction thus reducing reagent costs, sample preparation time and DNA requirements for these complementary analyses.

## Background

Bread wheat has a large complex allohexaploid genome that is 17GB in size and made up from three progenitor genomes (AABBDD). This size makes whole genome resequencing costly^1^. Therefore, a number of genome complexity reduction techniques exist that make re-sequencing cost effective. These include approaches such as: Restriction site Associated sequencing or RAD-seq^2^, involving digesting DNA with restriction enzymes and sequencing a tag for each resulting fragment; transcriptome sequencing, where we sequence cDNA generated from mRNA^3^; sequence capture, using DNA hybridization to capture specific genome sequences then sequencing the captured fragments. With a high-quality draft wheat genome now available, it is possible to design capture probe sets for tiling evenly across the genome and to use them to accurately genotype and anchor SNPs and CNVs to the genome^4^.

In addition to genetic variation, epigenetic variation also provides a source of natural variability contributing to changes in gene expression and phenotype. The most common form of DNA methylation is 5-methylcytosine, an epigenetic mark found throughout the genome of most eukaryotic organisms. Cytosine methylation has been implicated with orchestrating the structure and function of the genome, regulating chromatin and gene expression and it is found in plants in the context of CG, CHG and CHH^5,6^. It is thought that cytosine methylation maybe important for plants, providing a mechanism for rapidly adapting to environmental change.

Bisulphite treatment deaminates unmethylated cytosines resulting in conversion from a cytosine to a uracil residue. Therefore, bisulphite treatment in combination with sequencing can identify methylated cytosine residues, an approach termed methyl-seq^7^. Previously, we used methyl-seq in combination with sequence capture to survey the epigenome in hexaploid bread wheat^8^. An important question now is to understand how methylation varies across a globally diverse collection of wheat germplasm adapted to specific local agricultural niches. However, to apply methyl-seq to this kind of dataset you ideally require both DNA sequence data and bisulphite treated sequence data for each wheat accession otherwise C-T SNPS will be incorrectly classified as unmethylated cytosine sites.

Here, we describe a new wheat capture probe set that is tiled across the hexaploid bread wheat genome. We present a method, based on Agilent SureSelect Methyl-Seq, that will use a single capture assay as a starting point that is sequentially split and used for both DNA sequencing and methyl-seq. We benchmark the approach with the reference accession Chinese Spring and demonstrate its utility with an accession from the Watkins collection.

## Results

### Design of the wheat capture probe set (Supplementary Figure 1)

Probes were designed to capture a subset of wheat genes totaling 36Mb; 12Mb from each of the three sub-genomes of wheat. The design space was based on the 111Mb of assembled wheat genic sequence previously used for a NimbleGen (Roche) exome capture probe set. The wheat sequence had been processed to remove repetitive sequence, remove chloroplast and mitochondrial sequence, collapse redundant sequence and collapse homoeologous genes into one representative sequence (Gardiner *et al.*^4^ see Figure S1). Initially, 120bp sequences were tiled across the 111Mb of genic sequence at 40bp intervals resulting in 2.3 million potential probe sequences. These probes were then annotated with the following information: (i) % alignment to the International Wheat Genome Sequencing Consortium (IWGSC) reference sequence (positional information and gene annotations were also recorded); (ii) number of homoeologous and varietal SNPs (utilising IWGSC, CerealsDB and the wheat ancestral genomes), to allow discrimination between the wheat sub-genomes and to capture diversity, respectively; and (iii) average depth of coverage of the region obtained in previous sequence capture experiments using NimbleGen probes^4,9,10^. These annotations were used to rank the probes and the ‘best’ 100,000 were selected for the capture probe set. In addition to the genome wide tiling, for genes identified as associated with drought tolerance (Supplementary table 1^11,12,13,14^) and the NB-ARC conserved domains of nucleotide-binding site leucine-rich repeat (NBS-LRR) disease resistance genes^15^, 120-mer probes were tiled end-to-end across these key sequences to ensure that they were enriched effectively. Finally, a bias for even tiling of probes across the chromosomes was implemented, i.e. where possible there was one probe per assembled contig with additional bias for available surrounding sequence to facilitate effective mapping.

The 120-mer cRNA capture probe or ‘bait’ sequences were uploaded to Agilent eArray (online custom microarray design tool) to allow submission for manufacture. Bait ‘boosting’ was selected to permit excess unused design space (less than 1Mb in this case) to be filled with repeat sequences of baits predicted to perform less efficiently i.e. those with an above average GC content are ‘boosted’ to ultimately gain even depth of sequence coverage across the target region.

### Sequence capture

Using our custom probe set, we follow a SureSelect Methyl-Seq library preparation and hybridisation protocol, however we divide the sample immediately after capture and take one aliquot through Illumina paired-end sequencing protocol and the second aliquot through bisulphite conversion and Illumina paired-end sequencing. In order to assess the quality of the sequencing data generated we also performed standard SureSelect and SureSelect Methyl-Seq enrichments followed by sequencing and compared the output. This splitting after capture methodology allows us to directly compare the genotype and epi-type of the same DNA sample.

Total genomic DNA was extracted from 10-day old Chinese Spring wheat seedlings grown at 22°C. Three 3μg aliquots of the genomic DNA were fragmented to an average size of 200bp and following purification, each aliquot was used as input material in three separate SureSelect target enrichments. For the first enrichment, (i) a standard SureSelect library was prepared using 5 cycles of PCR to amplify the adapter-ligated DNA. 750ng of the pre-capture library was hybridised to 5μl of wheat custom baits. After post-capture washing, the enriched library was amplified (on-bead) and indexed using 10 PCR cycles; this is subsequently referred to as non-bisulphite treated full (NBTF). For the second enrichment, (ii) a SureSelect Methyl-Seq library was prepared and hybridised to our custom baits. Following post-hybridisation washing, the enriched DNA was eluted from the streptavidin beads, bisulphite treated and amplified using 8 cycles, followed by 6 indexing cycles as per the standard protocol; this is subsequently referred to as bisulphite treated full (BTF). For the third enrichment (Figure 1), (iii) a Sureselect Methyl-Seq library was again prepared and hybridised as usual but the enriched DNA was eluted from the streptavidin beads in a greater volume of SureSelect Elution Solution – 27μl instead of 20μl. At this point the eluted DNA was divided at a ratio of approximately 3:1. 20μl was taken into bisulphite treatment and amplification according to the standard protocol used for (ii); this is subsequently referred to as bisulphite treated split (BTS). The remaining 7μl of eluted DNA was neutralised by addition of an equal volume of SureSelect Neutralisation Solution, purified and then amplified alongside the bisulphite treated sample using the same SureSelect Methyl-Seq amplification mixture, primers and cycle numbers; this is subsequently referred to as non-bisulphite treated split (NBTS). Enriched libraries were sequenced on an Illumina HiSeq 2500 generating 2 × 125bp paired-end reads. Protocols are described in detail in the Methods section.

**Figure 1.**
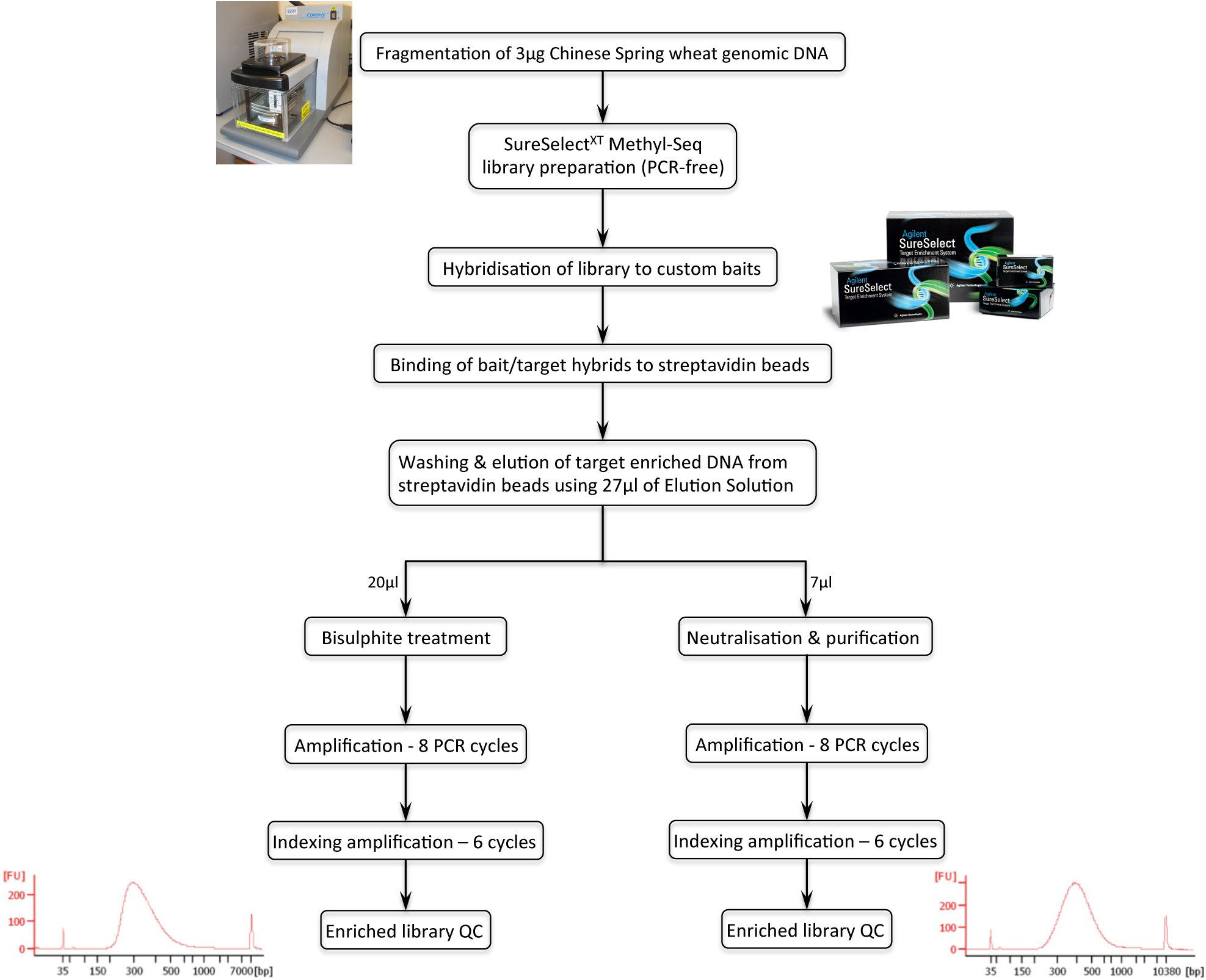
Workflow of the modified sequence capture method. Following fragmentation of the genomic DNA, a SureSelect Methyl-Seq library was constructed and hybridised to custom baits. Bait/target hybrids were bound to streptavidin beads, which were then washed to remove non-specifically bound DNA fragments. Target enriched DNA was eluted from the streptavidin beads and the eluate divided; ~¾ of the eluate was bisulphite converted and then amplified, ~¼ was neutralised, purified and then amplified. The quality of the purified libraries was assessed prior to sequencing.

### Our methodology has no detrimental effect on capture efficiency

Using paired-end sequencing reads to extend into the regions surrounding the capture probes, the mapped space exceeds the capture probe set design of 12Mb by more than 4x and 3x in the data from the non-bisulphite treated and bisulphite-treated samples, respectively. Looking at the non-bisulphite treated datasets, full and split enrichments were highly comparable with neither more than 1.2% from their average sequencing depth of 35.7x, across 51.7Mb of the extended reference bait sequence (Table 1). The depth of coverage across the probe set was summarised for the non-bisulphite treated samples across pseudo chromosomal molecules that were generated using POPseq data^16^ and coverage was relatively consistent with most falling into the range 5-70x (Supplementary Figure 2). On average, only 6% of baits exceeded 2x the average depth of coverage and ~4% showed an average coverage less than 5x. 97.4% of SNPs were conserved between the full and split samples at positions that were mapped to a minimum depth of 10x per sample (948,282). Similarly, the full and split bisulphite treated samples had an average depth of coverage of 30.6x, with neither more than 0.1% from this average, across 39.7Mb of the extended reference bait sequence.

**Table 1.** Mapping statistics for the reference sequence

| **Sample** | **Average %**<br>**coverage per ref**<br>**contig** | **Average depth**<br>**of coverage per**<br>**ref contig** | **Number of**<br>**ref contigs**<br>**mapped** | **% of ref**<br>**contigs**<br>**mapped** | **Base-space**<br>**mapped (bp)** |
| --- | --- | --- | --- | --- | --- |
| BTF | 59.6 | 30.7 | 82873 | 99.6 | 39,868,184 |
| BTS | 59.3 | 30.5 | 82862 | 99.6 | 39,602,695 |
| NBTF | 70.7 | 36.9 | 83107 | 99.9 | 48,808,952 |
| NBTS | 77.9 | 34.5 | 82999 | 99.7 | 54,641,687 |

Detailing the mapping output statistics for the two enriched wheat DNA samples, NBTF and BTF (non-bisulphite treated and bisulphite treated) that were taken through separate capture reactions and the two samples that were split and one bisulphite treated while the other was non-bisulphite treated after a single capture (NBTS and BTS). Mapping statistics are in relation to the 82.5Mb mapping reference

For the non-bisulphite treated datasets there were 49.4 million sequencing reads in the non-split sample; 73.3% of these reads could be mapped to our reference and 56.2% of these were identified as duplicate reads and removed leaving 32% of reads for analysis. In the split sample, there were 44.8 million sequencing reads; 72.8% could be mapped to our reference with 49.8% identified as duplicate reads leaving a highly comparable 36.6% of reads for analysis.

For the bisulphite treated datasets there were 49.9 million sequencing reads in the non-split sample; 24.2% of these reads could be mapped to our reference and only 10.1% of these were identified as duplicate reads and removed leaving 21.8% of reads for analysis. In the split sample, there were 50 million sequencing reads; 24.2% could be mapped to our reference with 11.3% identified as duplicate reads leaving again a highly comparable 21.5% of reads for analysis.

For the bisulphite treated full and split samples, differential methylation between the A, B and D sub-genomes was recorded using the tool methylKit to identify a minimum difference of 25% and p<0.01 (see Methods). 239,100 residues were available for comparison between the two samples i.e. residues with a depth of 5x or more per sub-genome in both samples. Of these 239,100 residues, only 0.006% showed methylation differences. This methylation between the samples is more similar than that seen between biological replicate samples in our previous studies (<0.09% difference observed^8^) and highlights our maintained capability to confidently define methylation patterns even after sample splitting. Furthermore, Pearson correlation plots demonstrate high comparability between samples with correlation coefficients consistently at 0.97 when sub-genomes are compared between the full and split datasets (Figure 2).

**Figure 2.**
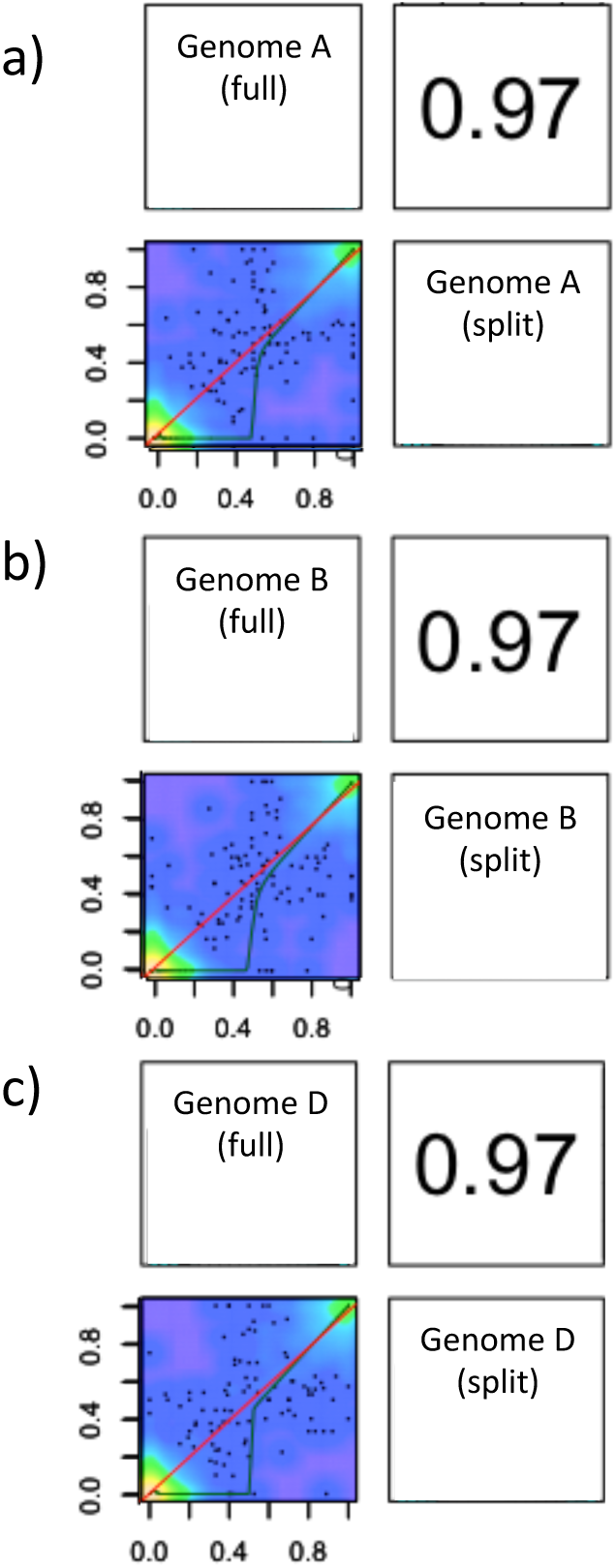
methylKit Pearson correlation coefficient computations to compare methylation between the split and non-split samples. Figures demonstrate comparisons of methylation levels across the respective bisulphite treated split and non-split samples at positions that have information associated with the a) A sub-genome, b) B sub-genome and the c) D sub genome. Individual samples are labelled diagonally through the middle of the plot, this diagonal axis acts as a mirror image division; comparative correlation plots lie to the left of the axis at the intersection between the two samples, with the corresponding correlation coefficient for the plot to the left of the axis at the intersection between the two samples.

Mapping the bisulphite treated sequencing reads to the non-methylated chloroplast genome was used to assess bisulphite conversion efficiency i.e. the percentage of cytosine bases that were successfully bisulphite converted^17,18^. While we did not enrich for chloroplast DNA specifically, a proportion of our off-target sequences mapped to the wheat chloroplast genome. Mapping statistics are shown in Table 2 where a consistently high level of coverage was gained (>350x) and highly comparable conversion efficiencies of 98.73% for the full sample and 98.82% for the split sample were observed.

**Table 2.**
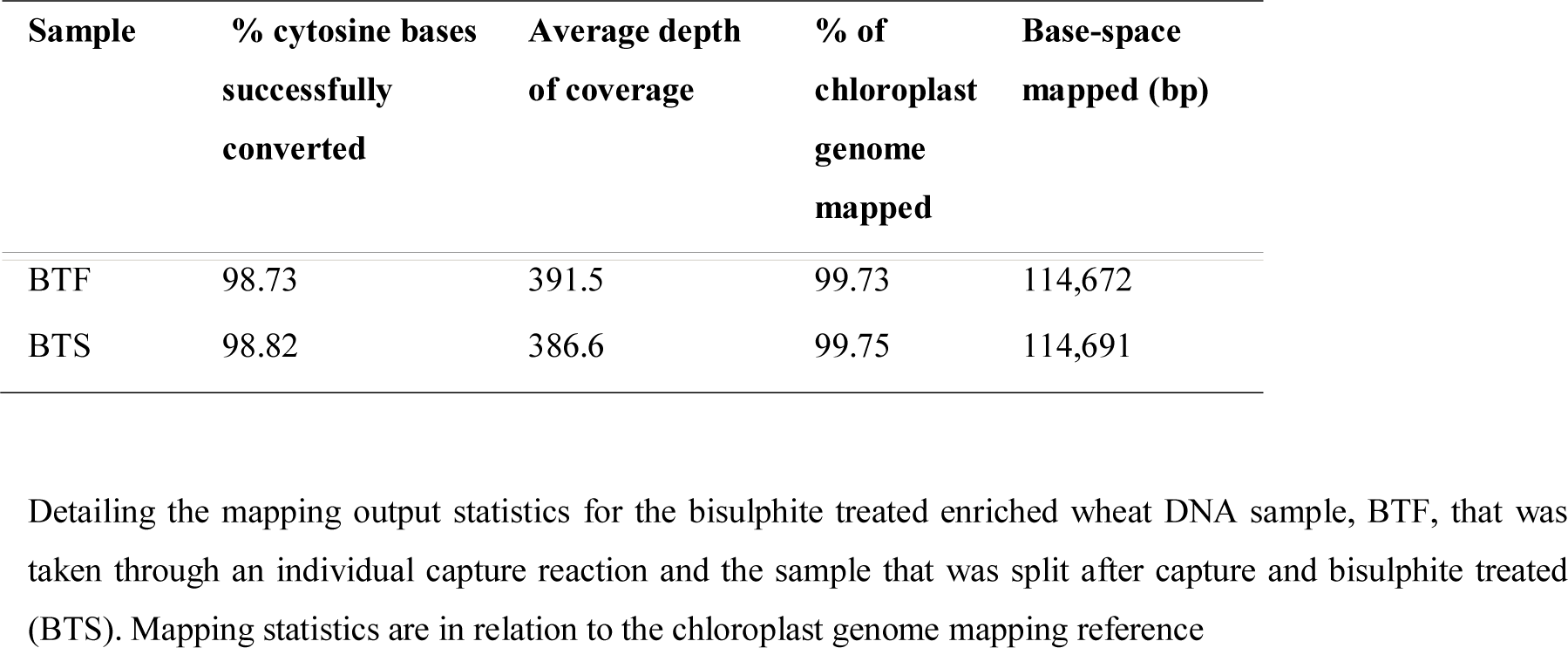
Mapping statistics for the chloroplast genome

### Demonstrating the utility of this method and capture probe set using a diverse wheat landrace

Our split after capture protocol was followed exactly as per for Chinese Spring to generate a bisulphite treated split (BTS) and non-bisulphite treated split (NBTS) library, however this time for a line from the Watkins bread wheat diversity collection that was chosen at random (line 1190103). Enriched libraries were sequenced on an Illumina HiSeq 4000 generating 2 × 125bp paired-end reads and sequencing reads were aligned to the mapping reference as per the methodology for Chinese Spring.

The split bisulphite treated sample had an average depth of coverage of 42.4x across 42.3 Mb of the extended reference bait sequence while the split non-bisulphite treated sample had an average depth of coverage of 66.7x across 51.1 Mb (Table 3). This is highly comparable to the mapping coverage generated by Chinese Spring (39-54 Mbp mapped), therefore the capture probe set can successfully enrich diverse wheat landraces that are thought to show a high SNP density compared to the reference accession Chinese Spring that is the basis of the capture design.

**Table 3.** Mapping statistics for the reference sequence (Watkins line 1190103)

| **Sample** | **Average %**<br>**coverage per ref**<br>**contig** | **Average depth**<br>**of coverage per**<br>**ref contig** | **Number of**<br>**ref contigs**<br>**mapped** | **% of ref**<br>**contigs**<br>**mapped** | **Base-space**<br>**mapped (bp)** |
| --- | --- | --- | --- | --- | --- |
| BTS | 64.8 | 42.4 | 81634 | 98.1 | 42,266,334 |
| NBTS | 76.3 | 66.7 | 82970 | 99.7 | 51,111,629 |

Detailing the mapping output statistics for the two Watkins wheat 1190103 samples that were split and one bisulphite treated while the other was non-bisulphite treated after a single capture (NBTS and BTS). Mapping statistics are in relation to the 82.5Mb mapping reference

SNPs were defined for the non-bisulphite treated dataset yielding 2,022,551 SNPs at a minimum of 10x. Of these SNPs, 672,949 were C→T or G→A SNPs that could be incorrectly classified as unmethylated cytosine sites if they were unidentified. Furthermore, 779,185 SNPs resulted in a C/G residue in the Watkins line where there was an A/T previously and these represent key missed opportunities where accession specific methylation, from accession specific cytosine residues that deviate from the reference sequence, may not have been previously analysed and therefore identified. The bisulphite treated sequencing data enables the analysis of 5,962,239 cytosines that show sequencing coverage at a minimum of 10x that is sufficient for accurate methylation calls; correction of the reference sequence for this Watkins line using the 672,949 C/G→T/A SNPs has the potential to eliminate up to 11.3% of these calls that were likely to be inaccurate and correction of the reference sequence using the 779,185 A/T→C/G SNPs would increase the cytosine set for analysis by approximately 1/5th.

This analysis demonstrates the utility of the capture probe set to enrich a diverse wheat accession that is likely to show a high SNP density compared to Chinese Spring while also quantifying the extent of the problems that we may encounter by not genotyping while we epi-type i.e. define how many residues could be given false positive methylation calls due to SNPs.

## Discussion

Here, we describe a new wheat capture probe set that is tiled across the hexaploid bread wheat genome. This capture probe set is evenly tiled across the genome and enriches typically over 4x the probe design space. It can be effectively utilised to survey and observe genome wide trends in wheat from a genotypic or epigenetic perspective. Furthermore, it can successfully enrich DNA from a diverse wheat accession from the Watkins landrace diversity collection despite it being designed based on the reference variety Chinese Spring.

Using this probe set, we present a method, based on Agilent SureSelect Methyl-Seq, that will use a single capture assay as a starting point that is sequentially split and used for both DNA sequencing and methyl-seq. This method is applicable to any organism of interest and therefore has a much wider usage potential than the use on wheat that we demonstrate here as an example. Understanding variation across populations is a common scientific question and we want to understand how methylation changes across a globally diverse collection of wheat germplasm using methyl-seq therefore we will require both DNA sequence data and bisulphite treated sequence data for each wheat accession otherwise CG-TA SNPS will be incorrectly classified as unmethylated cytosine sites. Furthermore, looking at a diverse landrace from the Watkins collection, this problem had the potential to affect 11.3% of sites, therefore this issue is of high priority to address. Moreover, correction of the reference sequence using A/T→C/G SNPs could increase the cytosine set for analysis by 1/5^th^ yielding further benefit to analyses.

With a single Agilent SureSelect capture reaction costing in excess of £500 (probe set plus capture reagents based on purchasing a set of 16), by utilising a single capture for both genotype and epigenetic analysis, we can cut these considerable costs. These savings are in addition to the reduction in staff labour costs associated with performing a lower number of capture reactions. Furthermore, there is an additional benefit to performing only one capture reaction to generate genotype and epigenetic information if the DNA quantity that is available for an individual sample is restricted.

Software has been developed to allow sample genotyping directly from bisulphite treated sequencing data and although this would reduce costs, removing the need for both DNA sequence and bisulphite treated sequence data, this method depends heavily on having sequence information for both DNA strands in the effort to discriminate C-T SNPs. Here, we profile methylation in only one strand of DNA, and as such this requirement for both strands would double the probe capture space and increase sequencing and probe costs. Furthermore, due to the high complexity of genotyping directly from the bisulphite treated sequencing data, such methodologies are highly error prone with reported false positive/false negative SNP calling rates ranging from 15% to upwards of 50%^19^. We, therefore present a gold standard methodology to ensure highly accurate SNP calls and following on from this high quality methylation calls.

## Conclusions

If we wish to accurately profile DNA methylation in diverse wheat lines, then it is advantageous to also generate genotype information. Here, we describe a new capture probe set that is tiled across the hexaploid bread wheat genome and can be effectively utilised to survey genome wide trends in wheat from a genotypic or epigenetic perspective. Furthermore, we present a cost-effective method for performing both DNA sequencing and methyl-seq from a single capture reaction thus significantly reducing reagent costs and DNA requirements.

## Methods

### Genomic DNA extraction and QC

Genomic DNA was extracted from the areal tissue of 10-day old Chinese Spring wheat seedlings grown at 22°C using a DNeasy Plant Mini kit (Qiagen) according to the manufacturer’s instructions. The DNA was quantified using a Qubit double-stranded DNA high sensitivity assay kit and Qubit fluorometer (Invitrogen). 100ng of DNA was analysed electrophoretically on a 1% agarose gel alongside HyperLadder 1kb (Bioline) to determine DNA integrity. This indicated that the extracted DNA was high molecular weight, with minimal degradation and no evidence of RNA contamination. DNA purity was assessed by obtaining the 260/280 and 260/230 absorbance ratios on a NanoDrop ND-1000 spectrophotometer.

### Genomic DNA fragmentation

Three 3μg aliquots of the same genomic DNA were each made up to a total volume of 130µl with 10mM Tris-HCl, pH 8.0. After mixing, each was transferred to a separate Covaris AFA microTUBE with pre-split snap-cap (Product number 520045) and sheared to an average size of approximately 200bp using a Covaris S2 focused-ultrasonicator (duty cycle 10%, intensity 5, 200 cycles per burst for 6 × 60s using frequency sweeping). The size distribution of the fragmented DNA was assessed with an Agilent 2100 Bioanalyser using a high sensitivity DNA chip. Each DNA aliquot was then purified using 1.4 × AxyPrep Mag PCR Clean-Up beads (Axygen) with two 70% ethanol washes (400μl) and elution in 50μl of nuclease-free water (Ambion). Each aliquot of purified fragmented DNA was used as input material for standard SureSelect library preparation or SureSelect Methyl-Seq library preparation as described below.

### Standard SureSelect target enrichment

A standard SureSelect library was constructed and hybridised essentially as described by the manufacturer in the SureSelect^XT^ Target Enrichment System for Illumina Multiplexed Sequencing Protocol; Version B.1, December 2014 (Agilent Manual Part Number G7530-90000), except all purification steps were carried out using AxyPrep Mag PCR Clean-Up beads instead of AMPure XP beads, since the former were more economical. Briefly, following end-repair, 3′-adenylation and paired-end adapter-ligation, 15μl (approximately half) of the adapter-ligated DNA was used as template in the pre-capture PCR with 5 cycles of amplification. The purified pre-capture library was quantified by Qubit double-stranded DNA high sensitivity assay and the quality assessed on a Bioanalyser high sensitivity DNA chip. The DNA fragment size peak was 245bp and the average fragment size was approximately 300bp.

Based upon the concentration obtained by Qubit quantification, 750ng of the pre-capture library was dehydrated until just dry by centrifugation under vacuum at 30°C in an Eppendorf Concentrator 5301. The DNA was then re-dissolved in 3.4μl of nuclease-free water and hybridised for approximately 20 hours at 65°C to 5μl of biotinylated custom SureSelect cRNA baits targeting the desired 12Mb of wheat sequence. The hybridisation was set up according to the Agilent protocol using 2μl of 25% RNase Block since the target was >3.0Mb. At the end of the hybridisation, bait/target hybrids were bound to 50μl of Dynabeads MyOne Streptavidin T1 magnetic beads (Invitrogen). Following post-capture washing, the target-enriched library was resuspended in 30μl of nuclease-free water and stored at −20°C for ~ 84h. Approximately half (14μl) of the bead-bound library was subsequently amplified with a primer containing an 8bp index using 10 PCR cycles. The purified captured library was quantified by Qubit double-stranded DNA high sensitivity assay and the size distribution ascertained by analysis on a Bioanalyser high sensitivity DNA chip. The library peaked at 287bp with an average fragment size of approximately 330bp.

### SureSelect Methyl-Seq Target Enrichment

A SureSelect Methyl-Seq library was prepared and hybridised by following the manufacturer’s instructions in the SureSelect^XT^ Methyl-Seq Target Enrichment System for Illumina Multiplexed Sequencing Protocol; Version C.0, January 2015 (Agilent Manual Part Number G7530-90002). The guide was followed from end-repair onwards, and again AMPure XP beads were replaced by AxyPrep Mag PCR Clean-Up beads. Quality assessment after end-repair was omitted since the DNA had already been analysed immediately after fragmentation.

Following methylated adapter ligation, the DNA was purified with elution in 25μl of nuclease-free water. The DNA size distribution was assessed on a Bioanalyser high sensitivity DNA chip and the library found to have a peak size of approximately 250bp and an average fragment size of 300bp. DNA concentration was determined by Qubit double-stranded DNA high sensitivity assay. The total yield of methylated adapter-ligated DNA was approximately 1.3μg and all of this was concentrated as previously described, reconstituted in 3.4μl of nuclease-free water and then used in the hybridisation step. The latter was conducted as described in the Agilent Manual but the Human methyl-seq capture library baits were substituted by 5μl of our 12Mb wheat-specific SureSelect baits. After approximately 20 hours at 65°C, bait/target hybrids were bound to streptavidin beads. Following post-capture washing, the bead-bound captured DNA was eluted with 20μl of SureSelect Elution Solution and bisulphite-treated using an EZ DNA Methylation-Gold Kit (Zymo Research) according to the instructions in the Agilent protocol. At this point, the bisulphite-converted and desulphonated library was stored at −20°C for ~ 84h. The library was then amplified using 8 cycles for the first PCR and 6 cycles for the indexing PCR with an indexing prime containing an 8bp index. The final library was quantified by Qubit double-stranded DNA high sensitivity assay and analysed electrophoretically on a Bioanalyser high sensitivity DNA chip. The library fragments had an average size of 360bp, with a peak at approximately 300bp.

### Modified SureSelect protocol

A SureSelect Methyl-Seq library was constructed and hybridised exactly as described in the previous section. Based on quantification obtained by Qubit double-stranded DNA high sensitivity assay, 1.2μg of methylated adapter-ligated DNA was obtained at the end of pre-capture library preparation. As previously, the DNA fragments peaked around 250bp and the average fragment size was 300bp when examined on a Bioanalyser high sensitivity DNA chip. The hybridisation was set up as outlined above using all of the pre-capture library as input. After the ~ 20 h 65°C hybridisation, bait/target hybrids were bound to streptavidin beads and standard post-capture washing was carried out. This time, 27μl of SureSelect Elution Solution was used to elute the target enriched DNA from the streptavidin beads. The beads were mixed with the Elution Solution and incubated at room temperature for 20 minutes, as instructed in the Agilent protocol. After this time, the beads and Elution Solution were separated using a DynaMag-2 magnet (ThermoFisher Scientific) and the supernatant divided into a 20μl aliquot and a 7μl aliquot – each being transferred to a separate 1.5ml Eppendorf tube. 7μl of SureSelect Neutralisation Solution was added to the 7μl aliquot of eluted DNA; after mixing by brief vortexing, the DNA was placed on ice. The 20μl aliquot of eluted DNA underwent bisulphite conversion and desulphonation according to the instructions in the Agilent protocol. During the bisulphite treatment (2.5h at 64°C, followed by 4°C hold), the other, neutralised aliquot was purified using 1.8 × AxyPrep Mag PCR Clean-Up beads. For this, 16μl of nuclease-free water was added to the 14μl mixture, bringing the volume up to 30μl. 54μl of AxyPrep Mag PCR Clean-Up beads were then added and a standard clean-up was carried out with two 70% ethanol washes (350μl) and elution with 19μl of nuclease-free water. At this point, the target-enriched, purified DNA (~ 19μl), and the enriched, bisulphite converted and desulphonated DNA (~ 20μl) were both frozen at −20°C for ~ 84h. The samples were then amplified in parallel according to the Agilent protocol. Although the two samples had been treated differently, the same amplification reagents and PCR cycling conditions were used. So, for the first PCR, each reaction contained 30μl of nuclease-free water, 50μl of SureSelect Methyl-Seq PCR Master Mix, 1μl of Methyl-Seq PCR1 Primer F, 1μl of Methyl-Seq PCR1 Primer R and 18μl of enriched DNA (bisulphite-treated or non-treated). The following cycling conditions were used: 95°C for 2 min, followed by 8 cycles of 95°C for 30 sec, 60°C for 30 sec and 72°C for 30 sec. A final extension of 72°C for 7 min was used followed by a hold at 4°C until further processing. The reactions were purified using 180μl of AxyPrep Mag PCR Clean-Up beads with two 70% ethanol washes (450μl) and elution with 21μl of nuclease-free water. Each eluate was then used as template in the final indexing amplification where each reaction contained 25μl of SureSelect Methyl-Seq PCR Master Mix, 0.5μl of SureSelect Methyl-Seq Indexing Primer Common, 5μl of Indexing Primer (containing an 8bp index) and 19.5μl of enriched amplified library (bisulphite-treated or non-treated). The cycling conditions were: 95°C for 2 min, followed by 6 cycles of 95°C for 30 sec, 60°C for 30 sec and 72°C for 30 sec. A final extension of 72°C for 7 min was used followed by a hold at 4°C. The reactions were purified using 90μl of AxyPrep Mag PCR Clean-Up beads with two 70% ethanol washes (450μl) and elution with 24μl of nuclease-free water. The final libraries were quantified by Qubit double-stranded DNA high sensitivity assay and analysed electrophoretically on a Bioanalyser high sensitivity DNA chip. The bisulphite-treated library peaked around 300bp with an average fragment size of 347bp. The non-treated library peaked at 390bp with an average size of 421bp.

### Illumina sequencing

All four of the libraries (two bisulphite-treated and two non-treated) were sequenced together with four other libraries of the same type. So, the eight libraries were pooled in equimolar amounts based on the Qubit and Bioanalyser data. The pool was further purified using 1.8 × AxyPrep Mag PCR Clean-Up beads. The size of the final pool was assessed on a Bioanalyser high sensitivity DNA chip and the DNA concentration was determined initially by Qubit double-stranded DNA high sensitivity assay, and then by qPCR, using an Illumina library quantification kit (KAPA) on a Roche LightCycler 480 II system. Sequencing was carried out on an Illumina HiSeq 2500, using version 4 chemistry, generating 2 × 125bp paired-end reads.

### Mapping reference sequence

The 12Mb of probe sequences align uniquely to 52,143 of the IWGSC reference gene contigs yielding partial representation of each. Utilising paired-end sequencing reads it is possible to extend the 120bp sequence that is captured by each probe to include surrounding regions. In previous studies this resulted in up to a 4x extension of coverage from the initial capture probe set. As such, in this case we anticipated capturing up to 48Mb per wheat sub-genome i.e. 144Mb overall. This necessitated a reference sequence that was constructed using the probes plus surrounding contiguous DNA sequence. These extended reference contigs ranged from 360bp-13168bp with a median length of 783bp. Therefore, in this study the total size of the mapping reference was ~82.5Mb per sub-genome.

### Standard mapping pipeline

All mapping analyses of non-bisulphite treated samples were carried out using BWAmem (version 0.7.10). Paired-end reads were mapped as fragment reads due to short reference contigs and only unique best mapping hits were taken forward^20^. Mapping results were processed using SAMtools; any non-uniquely mapping reads, unmapped reads, poor quality reads (<10) and duplicate reads were removed^21^. SNP calling in diploid datasets was carried out using the GATK Unified genotyper (after Indel realignment), which was used with a minimum quality of 50 and filtered using standard GATK recommended parameters, a minimum coverage of 5 and homozygous SNPs only were selected^22^. For polyploid datasets SAMtools mpileup was implemented with the SNP caller VarScan, to identify positions containing an alternate allele, with a minimum coverage of 5, an average mapping quality above 15 and a MAF of greater than 0.1^23^.

### Mapping of bisulphite treated DNA samples

The sequencing datasets for the samples were mapped to the extended probe sequence using Bismark, an aligner and methylation caller designed specifically for bisulphite treated sequence data. Sequencing reads were mapped as fragment reads rather than paired-end; a mismatch number of 3 was used and the non-directional nature of the library was specified^24^. The Bismark methylation extractor tool was then used to identify all cytosine residues within the mapping and categorize the reads mapping to them as un-methylated or methylated at that position while also detailing which type of potential methylation site was present (CHH, CHG or CpG). The mapping results were also processed for SNP calling using the standard polyploid pipeline described above.

### Determining a reference homoeologous SNP list

A reference homoeologous SNP list was determined across the 82.5Mb mapping reference using the same methods detailed by Gardiner *et al.*^8^. Firstly, the wheat ancestral genomes were aligned to the 82.5Mb reference to identify homoeologous SNPs directly. Secondly, non-bisulphite treated Chinese Spring sequencing reads were aligned to the IWGSC reference sequence to determine a genome of origin (only perfect and unique hits to one or two genomes were used). These genome assigned reads were then aligned to our single 82.5Mb reference sequence, which is representative of the 3 sub-genomes, allowing the discrimination of homoeologous SNP positions. SNP calling for polyploid datasets was carried out as previously described. Using genome assigned reads allowed us to match up the alleles at SNP locations with the contributing wheat sub-genome to define an additional homoeologous SNP list.

### Association of cytosine residues with the reference homoeologous SNP list

SNP positions were identified in the enriched hexaploid wheat bisulphite treated sequencing dataset using the standard polyploid pipeline. Reads mapping to these SNP positions therefore have sufficient depth and average mapping quality overall and one or more alternate allele present. Those positions that could also be found in the homoeologous SNP list were selected for further analysis i.e. homoeologous SNPs within the treated data. Any sequencing read with a mapping quality over 20, containing a cytosine residue methylation status calculated by Bismark, plus a homoeologous SNP allele, can be identified. Its SNP allele can be matched to a sub-genome therefore associating methylation status of that cytosine residue with a wheat sub-genome. For each cytosine position a summary of the number of reads hitting it for each sub-genome and whether or not these reads are methylated can be produced.

### Implementation of methylkit

The software methylKit^25^ was used to identify regions of differential methylation. Our summary of each cytosine position plus the number of reads hitting it for each sub-genome and whether or not these reads are methylated can be formatted and used directly as input for such analysis. Variation or differential methylation was recorded between the split and non-split bisulphite treated samples per sub-genome of wheat i.e. pairwise comparisons were between sub-genome A-A, B-B and D-D. Due to the use of pairwise comparisons the Fisher’s exact test was used to discriminate statistically significant differences (p < 0.01 and methylation difference of ≥ 50%).

### Construction of pseudo chromosomes from capture design contigs

We made use of 21 wheat chromosomal pseudomolecules that were created by organising and concatenating the IWGSC CSS assemblies using POPSEQ data^16^. BLASTN was used to place the extended probe sequences onto these chromosomal pseudomolecules (E-value cutoff 1e-5, minimum sequence identity 90 and minimum length of 100bp)^26^. Relative positions for the capture design contigs along the chromosomal pseudomolecules could then be used to order them into our POPSEQ based pseudo-chromosomes. We desired 7 POPSEQ based pseudo-chromosomes, as per our capture probe set, that were representative of the 21 wheat chromosomes. Therefore the order of the capture design contigs along genome B’s chromosomal pseudomolecules 1-7 was preferentially utilised since the greatest number of contigs could be aligned to these sequences and therefore included (83%).

## NOMENCLATURE

Bp: base pair
Mbp: million base pairs
SNP: Single nucleotide polymorphism

## Declarations

### Ethics approval and consent to participate

“Not applicable”

### Consent for publication

“Not applicable”

### Availability of data and material

All sequencing datasets are available (study PRJEB21533) from the European Nucleotide Archive (http://www.ebi.ac.uk/ena/data/view/PRJEB21533). The 12Mbp probe set is available from the corresponding author on request.

### Authors’ contributions

The methodology development, enrichment and Illumina sequencing library preparation were performed by L.O. qPCR and Illumina sequencing were performed by A.L. The bioinformatic analysis was performed by L.G. The project was designed by A.H. The project was planned and conducted by L.G, L.O, A. L, J.K., N.H. and A.H. The paper was written by L.G, L.O and A.H. with assistance from A.L. and J.K. NBS-LRR sequences for the capture design were provided by BS and BW. All authors read and approved the final manuscript.

## Acknowledgements

DNA sequence was generated by The University of Liverpool Centre for Genomic Research (United Kingdom).

## References

1. Clavijo, B. J. et al. An improved assembly and annotation of the allohexaploid wheat genome identifies complete families of agronomic genes and provides genomic evidence for chromosomal translocations, Genome Research, 2017;27(5):885–896

2. Baird, N. A. et al. Rapid SNP discovery and genetic mapping using sequenced RAD markers, PLoS ONE, 2008;3:e3376

3. De Wit, P. et al. SNP genotyping and population genomics from expressed sequences – current advances and future possibilities, Mol Ecol, 2015;24:2310–2323.

4. Gardiner, L. J. et al. Using genic sequence capture in combination with a syntenic pseudo genome to map a deletion mutant in a wheat species. Plant J, 2014;80:895–904.

5. Zhang, X. et al.. Genome-wide high-resolution mapping and functional analysis of DNA methylation in Arabidopsis, Cell, 2006;126:1189–201.

6. Cokus, S. J. et al.. Shotgun bisulphite sequencing of the Arabidopsis genome reveals DNA methylation patterning, Nature, 2008;452:215–9.

7. Darst, R. P. et al.. Bisulfite Sequencing of DNA. In: Current protocols in molecular biology edited by Frederick M. Ausubel et al. 2010: Unit–7.917.

8. Gardiner, L.-J. et al. A genome-wide survey of DNA methylation in hexaploid wheat. Genome biology, 2015;16:273.

9. The International Wheat Genome Sequencing Consortium (IWGSC). A chromosome-based draft sequence of the hexaploid bread wheat (Triticum aestivum) genome. Science, 2014;345(6194), 1251788.

10. Wilkinson, P. A. et al. CerealsDB 2.0: an integrated resource for plant breeders and scientists. BMC Bioinformatics, 2009;10:P5 13, 219.

11. Liang, Y. et al. Prediction of Drought-Resistant Genes in Arabidopsis thaliana using SVM-RFE, PLoS one, 2011;6(7):e21750

12. Uga, et al. Control of root system architecture by DEEPER ROOTING 1 increases rice yield under drought conditions, Nature Genetics, 2013;45: 1097–1102

13. Huang, D. et al. The relationship of drought-related gene expression in *Arabidopsis thaliana* to hormonal and environmental factors, Journal of experimental biology, 2008;59(11):2991–3007

14. Hu, H. and Xiong, L. Genetic engineering and breeding of drought-resistant crops, Annual review of plant biology, 2014;65:715–741

15. Steuernagel, B. et al. Rapid cloning of disease-resistance genes in plants using mutagenesis and sequence capture, Nature Biotechnology, 2016;34:652–655

16. Chapman, J. A. et al. A whole-genome shotgun approach for assembling and anchoring the hexaploid bread wheat genome. Genome biology, 2015;16(1): 26

17. Fojtová, M. et al. Cytosine methylation of plastid genome in higher plants. Fact or artefact? Plant Sci, 2001;160;585–593.

18. Lister, R. et al. Highly integrated single-base resolution maps of the epigenome in Arabidopsis. Cell, 2008;133:523–536.

19. Gao, S. et al., BS-SNPer: SNP calling in bisulfite-seq data. Bioinformatics, 2015;15;31(24):4006–4008

20. Li, H. & Durbin, R. Fast and accurate short read alignment with Burrows-Wheeler transform. Bioinformatics, 2009;25:1754–1760.

21. Li, H. et al. The Sequence Alignment/Map format and SAMtools. Bioinformatics, 2009;25:2078–2079.

22. McKenna, A. et al. The Genome Analysis Toolkit: A MapReduce framework for analyzing next-generation DNA sequencing data. Genome Res, 2010;20:1297–1303.

23. Koboldt, D. C. et al. VarScan 2: Somatic mutation and copy number alteration discovery in cancer by exome sequencing. Genome Res, 2012;22, 568.

24. Krueger, F. & Andrews, S. R. Bismark: a flexible aligner and methylation caller for Bisulfite-Seq applications. bioinformatics.oxfordjournals.org, 2011

25. Akalin, A. et al. methylKit: a comprehensive R package for the analysis of genome-wide DNA methylation profiles, Genome Biology, 2012;13:R87

26. Altschul, S. F., Gish, W., Miller, W., Myers, E. W. & Lipman, D. J. Basic local alignment search tool. J. Mol. Biol. 1990; 215 403–410.

